# Structure of the human frataxin-bound iron-sulfur cluster assembly complex provides insight into its activation mechanism

**DOI:** 10.1101/561795

**Authors:** Nicholas G. Fox, Xiaodi Yu, Xidong Feng, Henry J. Bailey, Alain Martelli, Joseph F. Nabhan, Claire Strain-Damerell, Christine Bulawa, Wyatt W. Yue, Seungil Han

**Author notes:** These authors contributed equally (Nicholas G. Fox, Xiaodi Yu). Correspondence should be addressed to Wyatt W. Yue, Seungil Han.

## Abstract

Iron-sulfur clusters (ISC) are essential in all life forms and carry out many crucial cellular functions. The core machinery for *de novo* ISC biosynthesis, located in the mitochondria matrix, is a five-protein complex containing the cysteine desulfurase NFS1 that is activated by frataxin (FXN), scaffold protein ISCU, accessory protein ISD11, and acyl-carrier protein ACP. Deficiency in FXN leads to the loss-of-function neurodegenerative disorder Friedreich’s ataxia (FRDA). Recently crystal structures depicting the inactive 3- and 4-way sub-complexes of the ISC biosynthesis machinery, lacking the key activator FXN, have been determined. Here, the 3.2 Å resolution cryo-electron microscopy structure of the FXN-bound active human complex, containing two copies of the NFS1-ISD11-ACP-ISCU-FXN hetero-pentamer, delineates for the first time in any organism the interactions of FXN with the component proteins. FXN binds at the interface of two NFS1 and one ISCU subunits, modifying the local environment of a bound zinc ion that would otherwise inhibit NFS1 activity in complexes without FXN. Our structure sheds light on how FXN facilitates ISC production through unlocking the zinc inhibition and stabilizing key loop conformations of NFS1 and ISCU at the protein-protein interfaces, and offers an explanation of how FRDA clinical mutations affect complex formation and FXN activation.

## Introduction

Iron-sulfur clusters (ISC) are inorganic cofactors essential in all life forms with common roles in electron transfer, radical generation, and structural support.^1^ In eukaryotes, the *de novo* ISC assembly machinery is located in the mitochondrial matrix and requires a core complex comprising the proteins NF<u>S</u>1, IS<u>D</u>11, <u>A</u>CP, and ISC<u>U</u> (SDAU).^1, 2^ The NFS1 cysteine desulfurase facilitates a pyridoxal 5’ phosphate (PLP) cofactor to generate the sulfane sulfur from L-cysteine, and deliver it to the ISCU scaffold protein.^3, 4^ The accessory protein ISD11/LYRM4 is unique in eukaryotes, and was shown to stabilize NFS1 and interact directly with the acyl carrier protein ACP/NDUFAB1.^5, 6^ ISCU utilizes three of its conserved cysteine residues (Cys69, Cys95, Cys138) to combine the sulfane sulfur from NFS1 with an iron source, resulting in ISC formation. ISCU then exploits the highly conserved ‘LLPVK’ motif for interaction with the chaperones, such as GRP75/HSCB,^7^ for the downstream delivery to apo-targets. Whereas electrons required for ISC assembly most likely involves mitochondrial ferredoxin/ferredoxin reductase combination,^8^ the iron source remains unclear.

An intronic GAA repeat of *FXN* gene, resulting in deficiency of the frataxin (FXN) protein, causes autosomal recessive Friedreich’s ataxia (FRDA).^9^ The *in vivo* loss of FXN results in oxidative stress due to iron accumulation in the mitochondria, rendering FRDA a fatal and debilitating condition with chelation therapy as the only mainstay treatment option. FXN is a key allosteric regulator of ISC assembly, and stimulates NFS1 activity by binding the SDAU complex to form the five-way active SDAUF complex.^10^–^12^ Zn^2+^ ion has been found to completely inhibit the SDAU complex *in vitro*, although its activity is restored by addition of FXN.^13^ Recently, crystal structures of SDA/SDAU/SDAU-Zn^2+^ complexes without the key component FXN have been published,^5, 14^ which attributed the zinc inhibition to the sequestration of key NFS1 catalytic residue Cys381, but could not serve as template to understand the molecular roles of FXN activator. To this end, we pursued structure determination of the SDAUF complex, coupled with FXN binding studies, to decipher the FXN-mediated activation mechanism.

## Results and Discussion

### Recombinant production and cryo-electron microscopy of the SDAUF complex

FXN binding to the SDAU complex is dynamic, yielding low-µM dissociation constants (*K*_*d*_) by bio-layer interferometry (BLI) (Supplementary Fig. 1a-c), hence presenting challenges to isolate the SDAUF complex intact with all 5 components in proper stoichiometry. Our several attempts to generate the SDAUF complex by reconstitution of individually expressed components (Supplementary Fig. 2a,b) did not fully incorporate FXN. To remedy this, we co-expressed in *E. coli* a plasmid containing His_6_-ISD11-NFS1-ISCU, with a plasmid containing His_6_-FXN (Supplementary Fig. 2c). This produced excess FXN, shifting equilibrium towards formation of a stable and active SDAUF complex comprising human SDUF co-purified with *E. coli* ACP (ACP_ec_). We attempted to make the 5-way all human complex, without *e coli* ACP, by inserting human ACP (NDUFAB1) into the second site of the vector containing His_6_-FXN. Upon co-expression with the plasmid containing His_6_-ISD11-NFS1-ISCU, we observed a heterogeneous complex containing an approximately equimolar mixture of the desired human and contaminating *E.coli* ACP (Supplementary Fig. 2d). Based on previous reports,^5^ and the functional conservation of human and *E. coli* ACP, we continued our experiments with a homogenous complex containing *E. coli* ACP with human SDUF (hereafter “SDAUF”). The as isolated complex could still be inhibited by Zn^2+^ due to the dissociation equilibrium of FXN (Supplementary Fig. 3, ‘SDAUF’), and the addition of more purified ISCU further exacerbated the Zn^2+^ inhibition (Supplementary Fig. 3, ‘SDAUF+U’). Zn^2+^ inhibition was fully reversed upon further FXN supplementation (Supplementary Fig. 3, ‘SDAUF+F’ and ‘SDAUF+U+F’), explaining the need for excess FXN to maintain its bound state within the five-way complex.

**Fig. 1.**
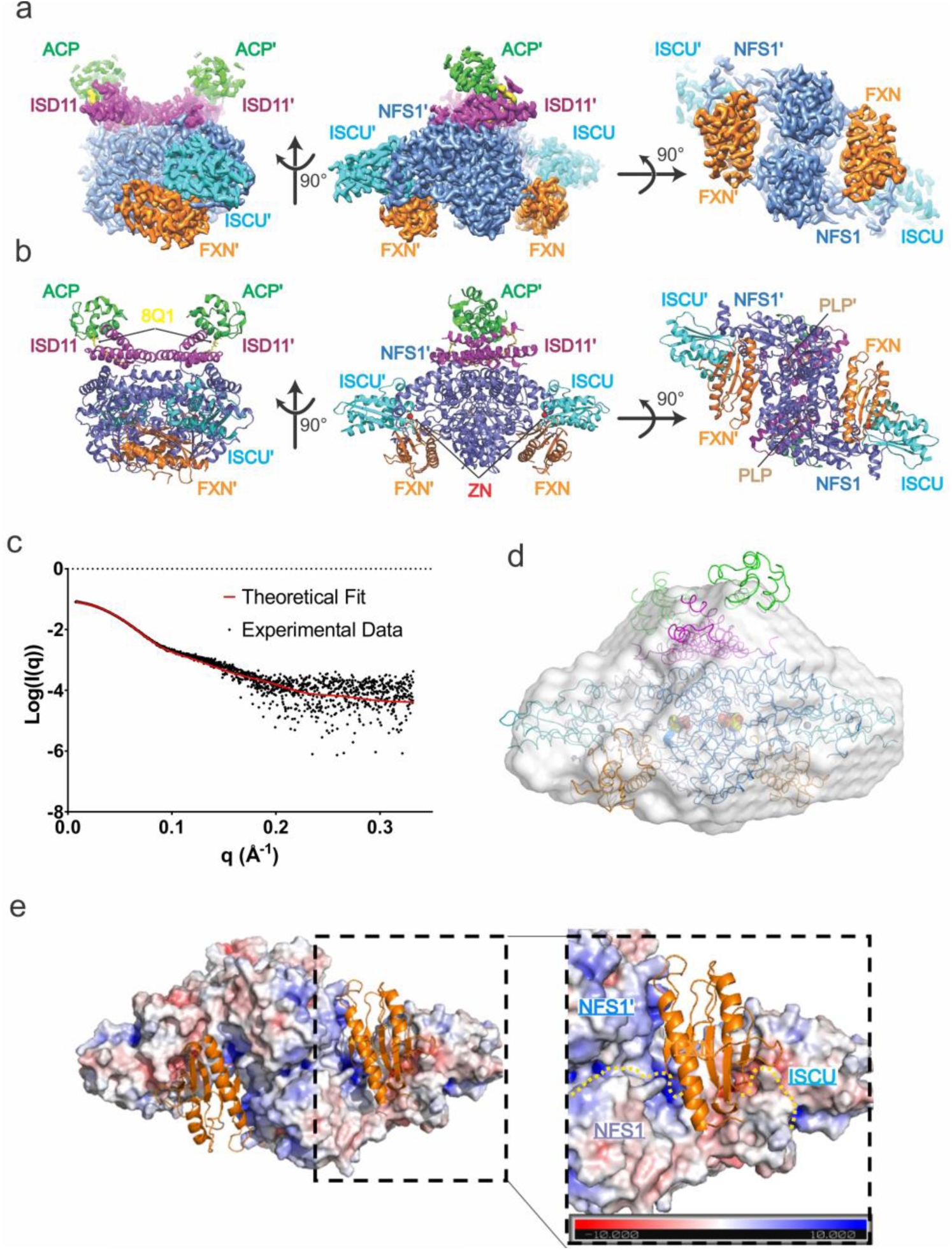
SDAUF-Zn^2+^ structure and FXN-NFS1 interactions. **a.** Cryo-EM density of SDAUF-Zn^2+^ structure (NFS1/NFS1’ slate, ISD11/ISD11’ magenta, ACP/ACP’ light green, ISCU/ISCU’, Cyan, FXN/FXN’ orange). **b.** Cartoon representation of SDAUF-Zn^2+^ complex. 4’-phosphopantetheine acyl chain (8Q1, yellow), Zn^2+^ ion (ZN, red) and pyridoxal 5’-phosphate (PLP, wheat) are shown as sticks/spheres. **c.** Scattering data from a sample of SDAUF used in cryo-EM was collected (black points) and fit to the theoretical SAXS profile back-calculated from the SDAUF-Zn^2+^ cryo-EM structure (red line) with χ^2^=1.98. **d**. The *ab initio* envelope calculated from SAXS data for the SDAUF sample was superimposed with the cryo-EM structure **e.** Each FXN (orange cartoon) binds to a cavity formed by interface (surface colored by electrostatic potential) of NFS1 homodimer and one ISCU. Yellow dotted lines denote protein boundaries.

**Fig. 2.**
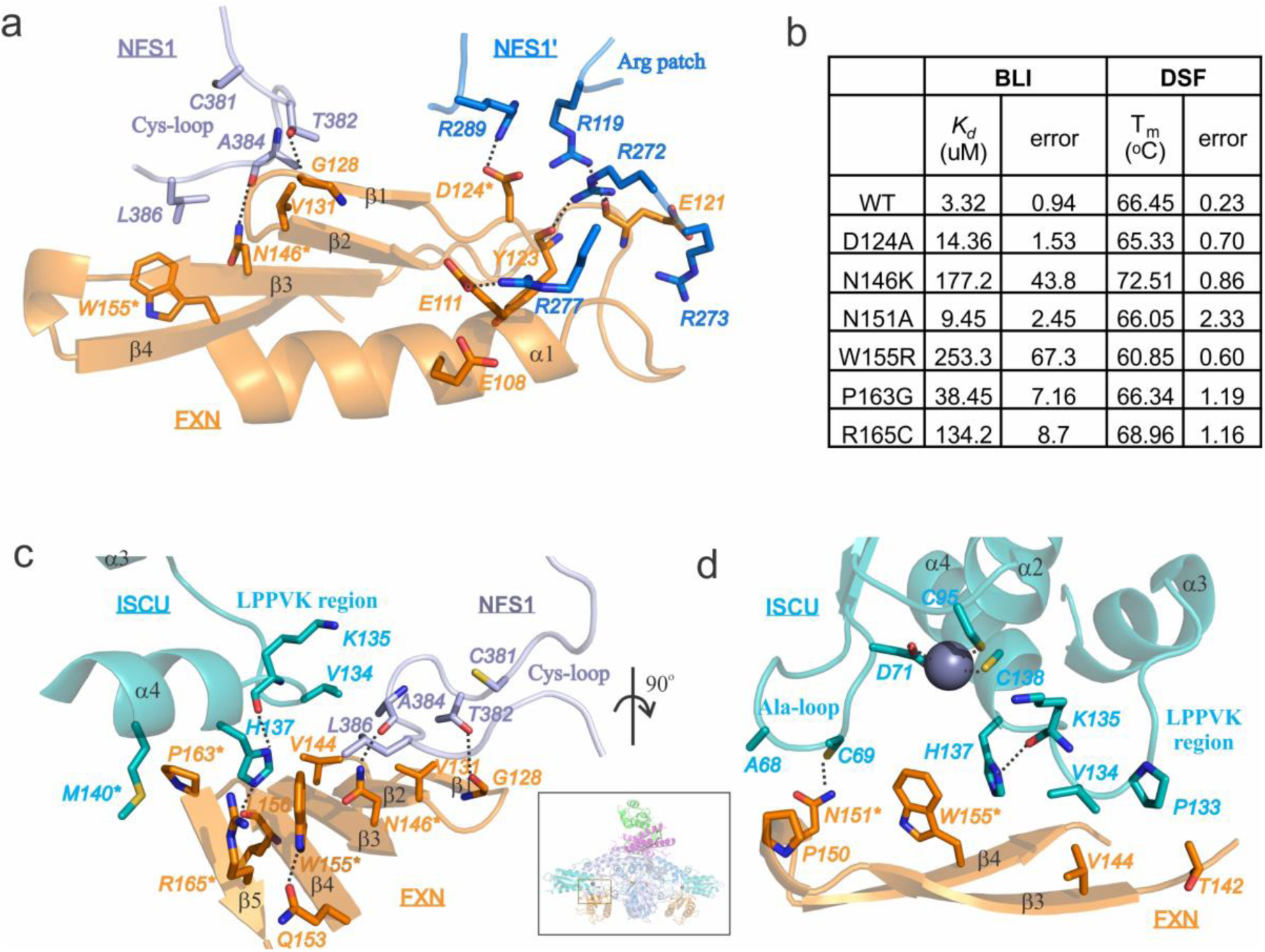
FXN-ISCU interactions and FXN mutagenesis. **a.** Interactions of FXN (orange) with both NFS1 subunits (NFS1, light slate; NFS1’, blue). Residues studied by site-directed mutagenesis are asterisked. Dashed lines denote hydrogen bonds. **b.** Determination of dissociation constant (*K*_d_, raw curves in Supplementary Fig. S1b) and melting temperature (T_m_) of FXN variants. **c.** Interface of FXN β-sheet (orange) with ISCU LPPVK-region (cyan) and NFS1 Cys-loop (slate). **d.** Interface of FXN with ISCU Ala-loop, LPPVK-region, and Zn^2+^ ion (sphere). *Inset*, viewpoints of panels c and d within SDAUF-Zn^2+^ complex.

**Fig. 3.**
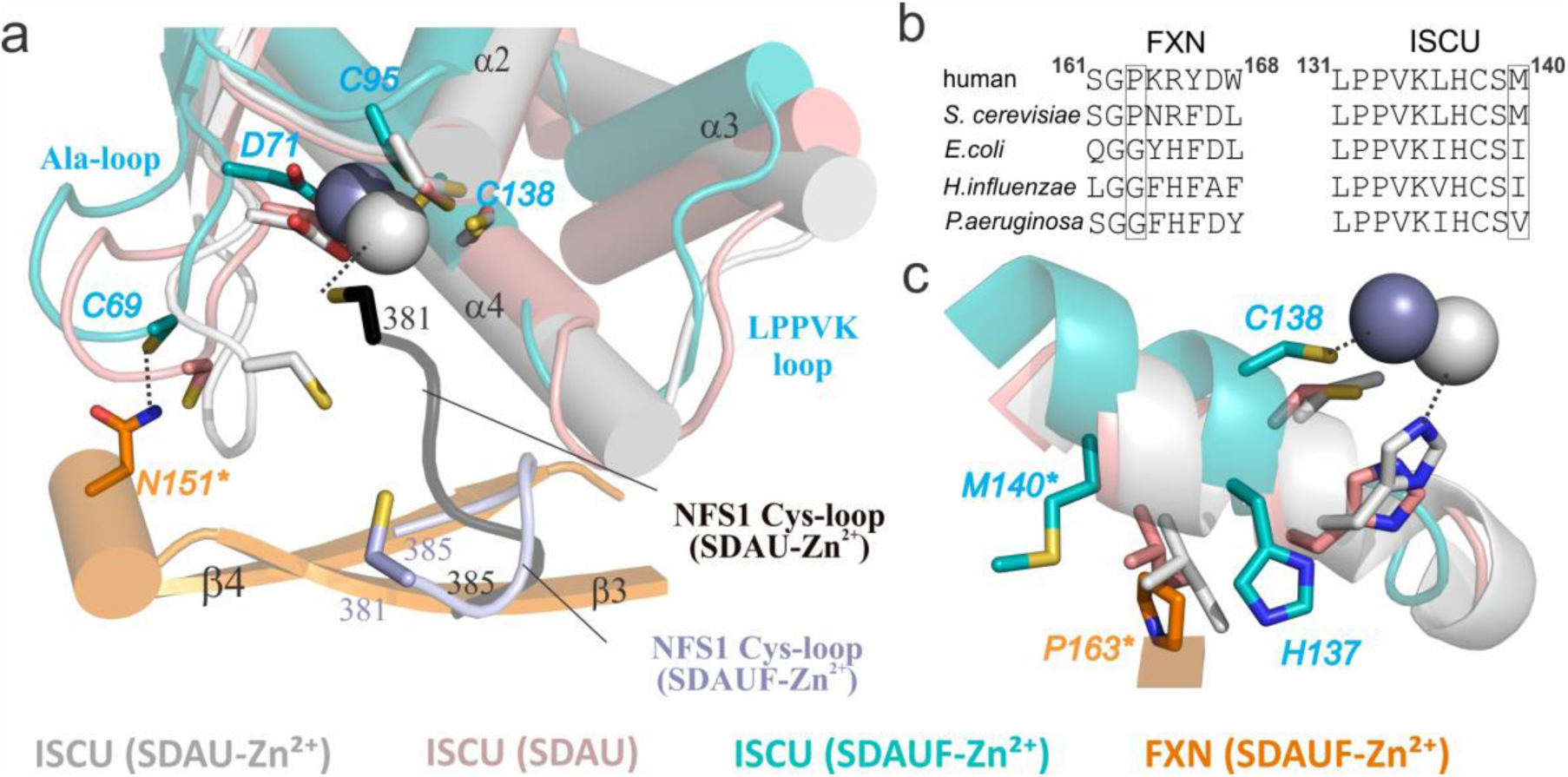
Comparing structures with and without FXN. **a.** Same view of SDAUF-Zn^2+^ as Fig. 2c, superimposed with ISCU subunits from SDAU-Zn^2+^ (5WLW), SDAU (5WKP) and *E. coli* IscS-IscU (3LVL) structures. NFS1 Cys-loops from SDAUF-Zn^2+^and SDAU-Zn^2+^ are shown. **b.** Sequence alignment of FXN region containing Pro163 (human numbering) and the ISCU L_131_PPVKLHCSM_140_ loop. Sequences shown include FXN and ISCU orthologues from human (Uniprot ID Q16595 and Q9H1K1, respectively), *S. cerevisiae* (Q07540, Q03020), *E. coli* (P27838, P0ACD4), *H. influenzae* (P71358, Q57074) and *P. aeruginosa* (Q9HTS5, Q9HXI9) **c.** ISCU L_131_PPVKLHCSM_140_ loop from the superimposed structures, adjacent to FXN Pro163. Human ISCU encodes Met at residue 140 (cyan), while in SDAU-Zn^2+^ and SDAU structures Met was substituted to Ile (white and pink respectively), the equivalent residue in *E. coli* IscS-IscU (purple). In both panels, FXN mutagenesis residues and ISCU Met140 are asterisked. Dashed lines denote hydrogen bonds or Zn^2+^ coordination.

We determined the single-particle cryo-electron microscopy (cryo-EM) structure of the 186-kDa SDAUF complex to 3.2 Å resolution (Supplementary Fig. 4 and Supplementary Table 1), allowing model building of entire complex components, with unambiguous placement of cofactor pyridoxal 5’-phosphate covalently-linked to NFS1 Lys258, a Zn^2+^ ion in each ISCU, and a 4’-phosphopantetheine (4’-PP) acyl-chain attached to ACP_ec_ Ser37 (Supplementary Fig. 5). LC-MS revealed a mixture of ACP components with different acyl-chains,^15^ and further top-down MS^3^ enabled the detailed structure elucidation on these acyl-chains through accurate mass measurements of both parent and fragment ions of the 4’-PP acyl chains which were readily ejected from ACP in the gas phase and interrogated further by tandem MS (Supplementary Fig. 6 and Supplementary Table 2). The relative abundances of the ACP components were estimated on the basis of extracted ion chromatograms in the LC-MS measurements, showing that ACP with longer acyl-chains are clearly the dominated species (Supplementary Table 2).

**Fig. 4.**
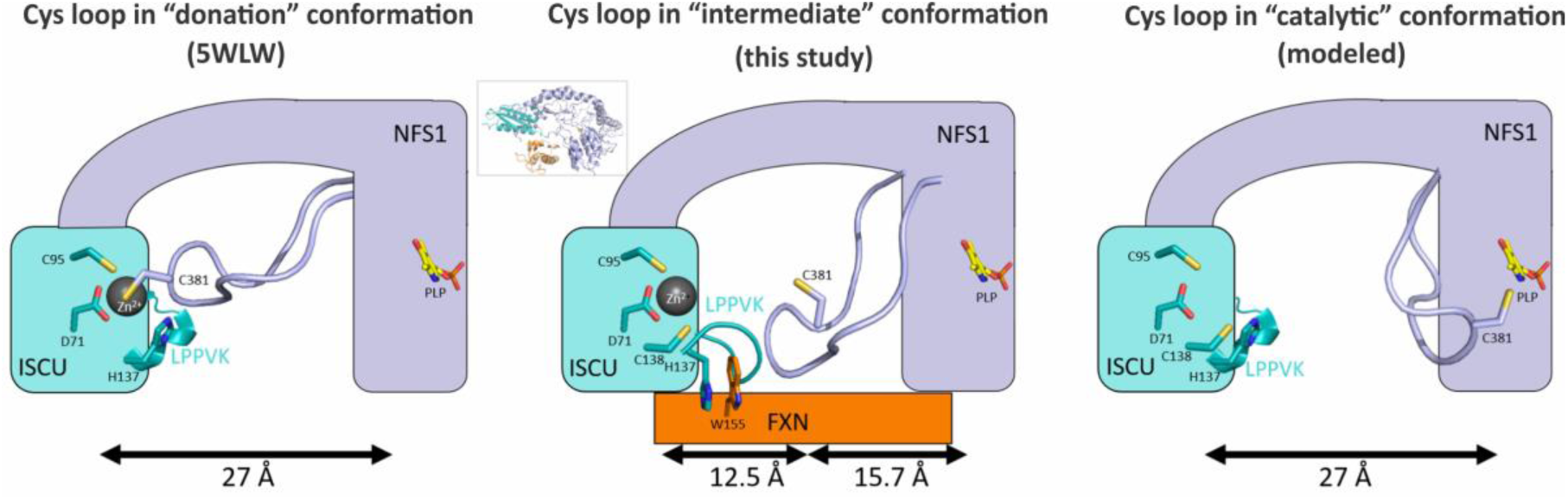
Schematic of the NFS1 Cys-loop trajectory, mediated by FXN-induced ISCU conformational changes. *Inset*, shows orientation of complex from which cartoon is derived.

### Overall architecture of the SDAUF complex

Our human SDAUF-Zn^2+^ structure, the first FXN-bound complex from any organism, is a symmetric heterodecamer comprising 2 copies each of the five proteins i.e. (NFS1)_2_(ISD11)_2_(ACP_*ec*_)_2_(ISCU-Zn^2+^)_2_(FXN)_2_. Structurally it constitutes a (NFS1-ISD11-ACP_*ec*_)_2_ homodimeric core, with one ISCU appended to each long end of the core, and one FXN fitted into the cavity next to each ISCU (Fig. 1a,b). This architecture agrees with small-angle x-ray scattering (SAXS) analysis (Fig. 1c,d and Supplementary Fig. 7a,b), whereby a theoretical SAXS profile back-calculated from our SDAUF-Zn^2+^ cryo-EM structure shows a good fit to experimental scattering data (χ^2^=1.98). While our five-way complex superimposes well with four-way SDAU/SDAU-Zn^2+^ structures from Boniecki *et al*^5^ *within the (NFS1-ISD11-ACP*_*ec*_)_2_ core (rmsd 0.6 Å), there is significant displacement of ISCU, up to 2.0 Å away from the core, in our FXN-bound complex (Supplementary Fig. 7c).

FXN occupies a cavity at the interface of both NFS1 and one ISCU subunits (Fig. 1e and Supplementary Fig. 8a), burying ∼1345 Å^2^ of FXN accessible surface. No direct FXN-ISD11 interaction is observed, contrasting previous predictions with oligomeric FXN.^16, 17^ A key feature of FXN binding is its simultaneous interactions with both NFS1 protomers of the complex (Fig. 2a), which definitively supports previous predictions from crosslinking, SAXS and NMR studies.^5, 18, 19^ Importantly, this requires a homodimeric arrangement of NFS1 within the complex, consistent with the SDAU conformation observed by Boniecki *et al*^5^, *while incompatible with the SDA*_*ec*_ *crystal structure from Cory et al* whereby an extensive NFS1 homodimer interface was not observed.^14^ The SDA conformation from Cory *et al*, unprecedented in all cysteine desulfurase structures published to date, would therefore not be conducive to the FXN activation mechanism, and could possibly instead depict a FXN-independent function.

### FXN Interactions with NFS1 dimer interface and C-terminus

FXN interacts with one NFS1 protomer *via* salt-bridges, and with the other NFS1 protomer through van der Waals contacts (Fig. 2a). The salt-bridge interface involves an acidic ridge of FXN (end of α1, loop α1-β1, start of β1) and a positively-charged Arg-rich patch (Arg272-Arg277) on one NFS1 protomer (Fig. 1d, 2a and Supplementary Fig. 8a,b). For example, FXN Asp124 can potentially form salt-bridges with NFS1 Arg289, while carbonyl backbones of FXN residues Glu121 and Tyr123 form H-bonds with Arg119 and Arg272 (Fig. 2a). The FXN(D124A) variant, aimed at abolishing these salt-bridges, binds SDAU with 4-fold weakened *K*_*d*_ compared to wild-type (WT)(Fig. 2b and Supplementary Fig. 1d). FXN contacts the other NFS1 protomer, and ISCU, using the β-sheet (β1-β5) surface (Fig. 2a,c,d). In this interface, FXN directly contacts the NFS1 loop containing catalytic residue Cys381 (‘Cys-loop’) (Fig. 2a), *via* hydrophobic interaction between FXN Trp155 and NFS1 Leu386, and H-bond between FXN Asn146 and NFS1 Ala384 carbonyl backbone. The clinical variant FXN(N146K) exhibited 50-fold weakened *K*_*d*_ towards SDAU (Fig. 2b and Supplementary Fig. 1d). Beyond the two extensive interfaces with FXN, NFS1 further contributes its C-terminal 20 aa (Ser437-His457) to wrap around the ISCU surface, with terminal residues Gln456-His457 anchored by FXN Asn151, Tyr175 and His177 *via* potential H-bonds (Supplementary Fig. 9). FXN(N151A) variant binds SDAU with 3-fold weakened *K*_*d*_ (Fig. 2b and Supplementary Fig. 1d). Therefore, NFS1, *via* two extensive interfaces and C-terminus, anchors FXN to interact with ISCU. This explains why FXN alone cannot bind ISCU without NFS1.^10, 12, 20^

### FXN binds to two key regions on ISCU

One ISCU-FXN interface is through the conserved ISCU Ala-loop (Ala66-Asp71), contributing the conserved Cys69 that is required for ISC biosynthesis and interacts with FXN Asn151 as well as Zn^2+^ coordinating ligand Asp71. This interaction, which may account for the weaker binding caused by the FXN(N151A) variant (Fig. 2b,d), is mediated by significant changes of the ISCU Ala-loop conformation in SDAUF as compared with SDAU-Zn^2+^ (rmsd ∼6Å), and zinc-free SDAU (rmsd ∼2Å) structures (Fig. 3a). The other, more predominant ISCU-FXN interface is through the conserved ISCU L_131_PPVKLHCSM_140_ sequence motif (Fig. 3b). This region (Fig. 2c,d), connecting ISCU helices α3 and α4, contains: the ‘L_131_PPVK_135_’ sequences recognized by the GRP75/HSCB chaperones for downstream ISC delivery,^7^ Cys138 the proposed sulfur acceptor for the NFS1 sulfane,^21^ and Met140, a residue reportedly determining if ISC biosynthesis is FXN-dependent (as in eukaryotes) or FXN-independent (prokaryotes).^22^

Previous structures of zinc-bound ISCU reveal a helical conformation for the L_131_PPVK_135_ region.^5^ In our structure, the displacement of ISCU caused by FXN (Supplementary Fig. 7d) is associated with the ISCU L_131_PPVK_135_ helix becoming loosened and more flexible, allowing His137 to pack against invariant FXN Trp155 (Fig. 2c,d). ISCU Pro133 and Val134 also pack against FXN Thr142 and Val144, respectively (Fig. 2d), in turn stabilizing NFS1 Cys-loop through interactions with NFS1 Ala384 and Leu386 (Fig. 2c). Considering the key role of FXN Trp155 in mediating ISCU and NFS1 conformations, the FXN(W155R) clinical variant of FRDA has >75-fold weakened *K*_*d*_ for SDAU compared to FXN(WT) (Fig. 2b and Supplementary Fig. 1d). Trp155 is further held in place by FXN Arg165 (pi-stacking) and Gln153 (H-bond) (Fig. 2c). The clinical variant FXN(R165C) has severely weakened *K*_*d*_ by 40-fold (Fig. 2b and Supplementary Fig. 1d).

Substitution of Met140 in yeast Isu, to the amino acids (Ile, Leu, or Val) observed at the equivalent prokaryotic position (Fig. 3b), obviated the need for Yfh1 (FXN equivalent) and reversed ΔYfh1 phenotype.^23^ Our structure shows that ISCU Met140 packs against FXN Pro163 (Fig. 3c), and a M140I substitution (present in the SDAU/SDAU-Zn^2+^ structures)^5^ sterically clashes with Pro163, unless the 163 position adopts the bacterial equivalent amino acid, Gly (*E. coli* Gly68; Fig. 3b). Our structure hence illustrates the evolutionarily distinct Met:Pro pairing in eukaryotes and Ile:Gly in prokaryotes. FXN(P163G) binds SDAU 10-fold weaker than FXN(WT) (Fig. 2b and Supplementary Fig. 1d).

### FXN modifies the Zn^2+^ environment and its influence within the complex

FXN binding to SDAU also influences the ISCU Zn^2+^ environment and NFS1 Cys-loop. In the reported SDAU-Zn^2+^ structure, ISCU Asp71, Cys95 and His137, and NFS1 Cys381 (from Cys-loop) form the Zn^2+^ ligation.^5^ This structure explained the Zn^2+^-dependent inhibition of NFS1 activity, due to sequestration of the catalytic Cys381 away from turning over the substrate cysteine. In our SDAUF-Zn^2+^ structure, zinc is ligated by ISCU Asp71, Cys95 and Cys138 (Fig. 2d). This rearranged metal coordination frees up ISCU His137 (now 3.7 Å away from Zn^2+^) to interact with FXN Trp155, Leu156 carbonyl backbone, Pro163, and ISCU Lys135 carbonyl backbone. Importantly, NFS1 Cys381 is also freed from Zn^2+^ ligation, now available for sulfur transfer (Fig. 3a). Our structure captured a novel conformation of NFS1 Cys-loop that positioned Cys381 approximately halfway between the NFS1 active site and conserved ISCU Cys residues, as part of a loop trajectory of 27 Å that could take place during ISC assembly (Fig. 4). Supported by the activity assay (Supplementary Fig. 3), our data therefore reveals how FXN unlocks the zinc inhibition of SDAU complex, to activate NFS1 into a conformation that is now poised for its incoming substrate cysteine.^13^

## Conclusion

To conclude, our structure elucidates how FXN binding to SDAU complex causes significant conformational changes to ISCU, which unlocks the zinc inhibition and primes its key regions (Ala-loop and LPPVK region) to facilitate mobility of NFS1 Cys-loop for sulfide formation and transfer during ISC assembly (Fig. 4). This work provides the framework for future mechanistic studies on the dynamics of SDAUF complex during next steps in ISC biosynthesis. For example, the ‘LPPVK’ region of ISCU, proposed to be important for downstream chaperone binding, is buried at the FXN interface in our structure, raising the questions whether FXN would be released from the complex to make way for binding ferredoxin/chaperones,^24^ and whether an ISC-loaded ISCU would be replaced by ‘apo-ISCU’ during the ISC biosynthesis cycle.

This work also represents one of very few reported cryo-EM structures of <200 kDa and >3.5 Å resolution for both membrane and soluble proteins. Our ability to visualize ligands and cofactors in such depth shows the advancement of modern cryo-EM for structure determination. We expect more examples to follow as the technique is applied to a broad range of clinically-relevant targets that remain intractable for x-ray crystallography.

## Methods

### Cloning, expression and purification of human ISCU, FXN, and NFS1-ISD11

For bi-cistronic co-expression of NFS1-ISD11 and tri-cistronic co-expression of NFS1-ISD11-ISCU, a DNA fragment encoding His-tagged ISD11, non-tagged NFS1 (Δ1-55), and for tri-cistronic additional non-tagged ISCU (Δ1-34) separated by an in-frame ribosomal binding site, was sub-cloned into pNIC28-Bsa4 vector. Constructs of human FXN (Δ1-80) was subcloned into the pCDF-LIC-Bsa4 vector and the construct of human ISCU (Δ1-34) isoform 1 was subcloned into the pNIC28-Bsa4 vector (GenBank ID: EF198106) for recombinant *E. coli* expression using BL21(DE3)-R3-pRARE2 cells. Cells transformed with the above plasmids were grown in Terrific Broth and induced with 0.1 mM isopropyl β-D-1-thiogalactopyranoside (IPTG) for 16 hours at 18 °C. Cell pellets were resuspended in binding buffer (50 mM HEPES pH 7.5, 500 mM NaCl, 20 mM Imidazole, 5% glycerol, and 2 mM TCEP) containing EDTA-free protease inhibitor (Merck), and lysed by sonication. For NFS1-ISD11 purification, 150 μM pyridoxal 5’-phosphate (PLP) was supplemented to the binding buffer during sonication.

The clarified supernatant was incubated with 2.5 mL Ni Sepharose 6 fast flow resin (GE Healthcare), washed and eluted with binding buffer containing 40 mM and 250 mM Imidazole, respectively. Elution fractions were collected, 10 mM DTT was added to samples containing ISCU or NFS1, and loaded onto gel filtration (Superdex S75 for ISCU and FXN, Superdex S200 for NFS1-ISD11 complex; GE Healthcare). Peak fractions were collected, treated with His-TEV protease to remove His-tag, and then passed onto Ni Sepharose 6 fast flow resin to remove His-TEV and cleaved His-tag. Fractions containing target protein were collected and buffer exchanged into gel filtration buffer (50 mM HEPES pH 7.5, 200 mM NaCl, 5% glycerol, and 2 mM TCEP) or cryo-EM buffer (20 mM HEPES pH 7.5, 100 mM NaCl, and 2 mM TCEP). As reported by others, recombinantly expressed NFS1-ISD11 complex co-purified with *E. coli* ACP, and will hereafter be referred to as the NFS1-ISD11-ACP (SDA) complex.

FXN variants were constructed using the Q5^®^ Site-Directed Mutagenesis Kit (NEB) and confirmed by sequencing of plasmid DNA and intact mass spectrometry of purified proteins. Cells transformed with the above plasmids were grown in 100 mL auto-induction Terrific Broth at 37 °C for 5 hours and then 20 °C for 2 days. Cell pellets were resuspended in Lysis Buffer (50 mM HEPES, pH 7.5, 500 mM NaCl, 5 % Glycerol, 2 mM TCEP, 1 µg/mL Benzonase, 1:1000 EDTA-free protease inhibitor (Merck), and 5 mg/mL lysozyme), left at room temperature for 30 min and then added 1 mL/g cells 10 % Triton X-100 and frozen for >1hour. Purification was similar to above but gel filtration was replaced with a PD-10 desalting column. All variants and wild-type were checked for stability using differential scanning fluorimetry and values recorded in Fig. 2e.

### Differential scanning fluorimetry

DSF was performed in a 96-well plate using an Mx3005p RT-PCR machine (Stratagene). Each well (20 µl) consisted of protein (2 µM in buffer containing 50 mM HEPES, pH 7.5, 250 mM NaCl, 5% glycerol, and 2 mM TCEP), SYPRO-Orange (Invitrogen, diluted 1000-fold of the manufacturer’s stock). Fluorescence intensities were measured from 25 to 96 °C with a ramp rate of 1 °C/min. *T*_m_ was determined by plotting the intensity as a function of temperature and fitting the curve to a Boltzmann equation. Final graphs were generated using GraphPad Prism. Assays were carried out in triplicate.

### Methylene blue activity assay

Sulfide production, due to cysteine desulfurase enzyme activity, was measured using the methylene blue colorimetric assay as described previously.^25, 26^ The standard assay was performed in buffer consisting of 50 mM HEPES pH 7.5, 200 mM NaCl, 10 mM DTT and either 100 μM EDTA or 10 μM ZnCl_2_. When noted, concentration of NFS1-ISD11-ACP (SDA) was at 0.5 μM, ISCU (U) at 2.5 μM, Frataxin (F) at 5 μM and 0.5 μM SDAUF were used and mixed in a 1.5 mL black Eppendorf tube with total volume of 800 μL. The reaction was initiated by adding 100 μM L-Cysteine and placed in 37 °C incubator for 10 min (with FXN) or 20 min (without FXN) and then quenched with 100 μL of 30 mM FeCl_3_ in 1.2 N HCl and 100 μL of 20 mM *N,N*-dimethyl-*p*-phenylenediamine (DMPD) in 7.2 N HCl and placed back in 37 °C incubator for 20 min, followed by centrifugation to spin down precipitant and then take the absorbance at 670 nm. Concentration of sulfide was calculated *via* a standard curve of Na_2_S.

### Bio-Layer Interferometry (BLI)

BLI experiments were performed on a 16-channel ForteBio Octet RED384 instrument at 25 °C, in buffer containing 50 mM HEPES pH 7.5, 200 mM NaCl, 5% Glycerol, 2 mM TCEP, 5 mM DTT, 0.5 mg/mL BSA. Biotinylated WT ISCU (U_b_) was combined with a purified batch of SDA and separated on Superdex 200 increase size exclusion column for isolation of the SDAU_b_ complexes. Complexes were at 0.1 mg/mL and loaded to the streptavin coated sensors. The concentration for FXN used ranged from 500 mM to 1 nM. Measurements were performed using a 90 second association step followed by a 60 second dissociation step on a black 384-well plate with tilted bottom (ForteBio). The baseline was stabilized for 30 sec prior to association and signal from the reference sensors was subtracted. A plot of response vs. [FXN] was used for *K*_*d*_ determination using one site-specific binding fit in GraphPad Prism (GraphPad Software).

### Small angle X-ray scattering (SAXS)

SAXS experiments for the SDAU and SDAUF complex were performed at 0.99 Å wavelength Diamond Light Source at beamline B21 coupled to the Shodex KW403-4F size exclusion column (Harwell, UK) and equipped with Pilatus 2M two-dimensional detector at 4.014 m distance from the sample, 0.005 < q < 0.4 Å^-1^ (q = 4π sin θ/λ, 2θ is the scattering angle). The samples were in a buffer containing 300 mM NaCl, 25 mM Hepes 7.5, 1 mM TCEP 2 % Glycerol, 1% Sucrose and the measurements were performed at 20°C. The data were processed and analyzed with Scatter and the ATSAS program package.^27^ Scatter was used to calculate the radius of gyration Rg and forward scattering I(0) via Guinier approximation and to derive the maximum particle dimension Dmax and P(r) function. The *ab initio* model was derived using DAMMIF^28^. 20 individual models were created, then overlaid and averaged using DAMAVER.^29^ FoxS^30, 31^ server was used for comparison of theoretical and experimental data.

### Grid preparation and data acquisition

3.5 µL of 1.5 mg/ml purified SDAUF complex was applied to the glow-discharged Quantifoil Au R1.2/1.3 grid (Structure Probe), and subsequently vitrified using a Vitrobot Mark IV (FEI Company). In order to overcome an orientation bias, n-octyl-β-d-glucopyranoside (BOG, Anatrace) was added to the sample prior freezing. Cryo grids were loaded into a Titan Krios transmission electron microscope (ThermoFisher Scientific) operating at 300 keV with a Gatan K2 Summit direct electron detector. Images were recorded with SerialEM in super-resolution mode with a super resolution pixel size of 0.543 Å and a defocus range of 1.2 to 2.5 μm. Data were collected with a dose rate of 5 electrons per physical pixel per second, and images were recorded with a 10s exposure and 250 ms subframes (40 total frames) corresponding to a total dose of 42 electrons per Å^2^. All details corresponding to individual datasets are summarized in Supplementary Table S1.

### Electron microscopy data processing

A total of 4,260 dose-fractioned movies were gain-corrected, 2 x binned (resulting in a pixel size of 1.086 Å), and beam-induced motion correction using MotionCor2 ^32^ with the dose-weighting option. The SDAUF particles were automatically picked from the dose-weighted, motion corrected average images using Gautomatch. CTF parameters were determined by Gctf. ^33^ A total of 1,316,416 particles were then extracted using Relion 2.0 ^34^ with a box size of 200 pixels. The 2D, 3D classification and refinement were performed with Relion 2.0. Two rounds of 2D classification and one round of 3D classification were performed to select the homogenous particles. After selecting particle coordinates, per-particle CTF estimation was refined using the program Gctf. ^33^ One set of 267,153 particles was then submitted to 3D auto-refinement with C2 symmetry imposed and resulted in a 3.2 Å map (Supplementary Fig. S4, S5). All 3D classifications and 3D refinements were started from a 60 Å low-pass filtered version of an ab initio map generated with VIPER. ^35^ To evaluate the contribution of imposed symmetry in the result, 3D refinement was repeated using the same set of 267,153 particles without imposing symmetry and produced a 3.4 Å map (Supplementary Fig. S4). Since the overall structures with/without imposing symmetry are nearly identical, the C2 symmetry density map was used for model building. All resolutions were estimated by applying a soft mask around the protein complex density and based on the gold-standard (two halves of data refined independently) FSC = 0.143 criterion. Prior to visualization, all density maps were sharpened by applying different negative temperature factors using automated procedures, ^36^ along with the half maps, were used for model building. Local resolution was determined using ResMap ^37^ (Supplementary Fig. S4).

### Model building and refinement

The initial template of the SDAUF complex was derived from a homology-based model calculated by SWISS-MODEL. ^38^ Each subunit was docked into the C2 symmetry EM density map using Chimera ^39^ and followed by manually adjustment using COOT. ^40^ The model was independently subjected to global refinement and minimization in real space using the module phenix.real_space_refine in PHENIX ^41^ against separate EM half-maps with default parameters. The model was refined into a working half-map, and improvement of the model was monitored using the free half map. The geometry parameters of the final models were validated in Coot and using MolProbity and EMRinger. ^42^ These refinements were performed iteratively until no further improvements were observed. The final refinement statistics were provided in Supplementary Table S1. Model overfitting was evaluated through its refinement against one cryo-EM half map. FSC curves were calculated between the resulting model and the working half map as well as between the resulting model and the free half and full maps for cross-validation (Supplementary Fig. S1). Figures were produced using PyMOL ^43^ and Chimera. ^39^

## ACCESSION CODE

All the cryo-EM data were deposited to the Protein Data Bank (PDB ID: xxx) and the EMDB (EMD-xxxx, EMD-xxxx).

## AUTHOR CONTRIBUTIONS

C.B., S.H., W.W.Y. designed the experiments. N.G.F., X.Y., X.F., H.J.B., A.M., J.F.N. and C.S-D. performed the experiments. N.G.F., S.H., and W.W.Y. wrote the manuscript.

## ACKNOWLEDGEMENTS

The Structural Genomics Consortium is a registered charity (Number 1097737) that receives funds from AbbVie, Bayer Pharma AG, Boehringer Ingelheim, Canada Foundation for Innovation, Eshelman Institute for Innovation, Genome Canada, Innovative Medicines Initiative (EU/EFPIA) [ULTRA-DD grant no. 115766], Janssen, Merck & Co., Novartis Pharma AG, Ontario Ministry of Economic Development and Innovation, Pfizer, São Paulo Research Foundation-FAPESP, Takeda, and Wellcome Trust [092809/Z/10/Z]. N.G.F. and W.W.Y. are further supported by funding from the Pfizer Rare Disease Consortium.

## REFERENCES

1 Lill, R. Function and biogenesis of iron-sulphur proteins. Nature 460, 831–838, doi:10.1038/nature08301 (2009).

2 Lill, R. & Muhlenhoff, U. Iron-sulfur protein biogenesis in eukaryotes: components and mechanisms. Annu Rev Cell Dev Biol 22, 457–486, doi:10.1146/annurev.cellbio.22.010305.104538 (2006).

3 Behshad, E. & Bollinger, J. M., Jr. Kinetic analysis of cysteine desulfurase CD0387 from Synechocystis sp. PCC 6803: formation of the persulfide intermediate. Biochemistry 48, 12014–12023, doi:10.1021/bi802161u (2009).

4 Zheng, L., White, R. H., Cash, V. L. & Dean, D. R. Mechanism for the desulfurization of L-cysteine catalyzed by the nifS gene product. Biochemistry 33, 4714–4720 (1994).

5 Boniecki, M. T., Freibert, S. A., Muhlenhoff, U., Lill, R. & Cygler, M. Structure and functional dynamics of the mitochondrial Fe/S cluster synthesis complex. Nat Commun 8, 1287, doi:10.1038/s41467-017-01497-1 (2017).

6 Van Vranken, J. G. et al. ACP Acylation Is an Acetyl-CoA-Dependent Modification Required for Electron Transport Chain Assembly. Mol Cell 71, 567–580 e564, doi:10.1016/j.molcel.2018.06.039 (2018).

7 Luo, W. I., Dizin, E., Yoon, T. & Cowan, J. A. Kinetic and structural characterization of human mortalin. Protein Expr Purif 72, 75–81, doi:10.1016/j.pep.2010.02.003 (2010).

8 Webert, H. et al. Functional reconstitution of mitochondrial Fe/S cluster synthesis on Isu1 reveals the involvement of ferredoxin. Nat Commun 5, 5013, doi:10.1038/ncomms6013 (2014).

9 Schmucker, S. & Puccio, H. Understanding the molecular mechanisms of Friedreich’s ataxia to develop therapeutic approaches. Hum Mol Genet 19, R103–110, doi:10.1093/hmg/ddq165 (2010).

10 Schmucker, S. et al. Mammalian frataxin: an essential function for cellular viability through an interaction with a preformed ISCU/NFS1/ISD11 iron-sulfur assembly complex. PLoS One 6, e16199, doi:10.1371/journal.pone.0016199 (2011).

11 Colin, F. et al. Mammalian frataxin controls sulfur production and iron entry during de Novo Fe4S4 cluster assembly. J Am Chem Soc 135, 733–740, doi:Doi 10.1021/Ja308736e (2013).

12 Tsai, C. L. & Barondeau, D. P. Human frataxin is an allosteric switch that activates the Fe-S cluster biosynthetic complex. Biochemistry 49, 9132–9139, doi:10.1021/bi1013062 (2010).

13 Fox, N. G. et al. Zinc(II) binding on human wild-type ISCU and Met140 variants modulates NFS1 desulfurase activity. Biochimie 152, 211–218, doi:10.1016/j.biochi.2018.07.012 (2018).

14 Cory, S. A. et al. Structure of human Fe-S assembly subcomplex reveals unexpected cysteine desulfurase architecture and acyl-ACP-ISD11 interactions. Proc Natl Acad Sci U S A 114, E5325–E5334, doi:10.1073/pnas.1702849114 (2017).

15 Majmudar, J. D. et al. 4’-Phosphopantetheine and Long Acyl Chain-Dependent Interactions Are Integral to Human Mitochondrial Acyl Carrier Protein Function. MedChemComm (2019 (in Press)).

16 Li, H., Gakh, O., Smith, D. Y. t. & Isaya, G. Oligomeric yeast frataxin drives assembly of core machinery for mitochondrial iron-sulfur cluster synthesis. J Biol Chem 284, 21971–21980, doi:10.1074/jbc.M109.011197 (2009).

17 Shan, Y., Napoli, E. & Cortopassi, G. Mitochondrial frataxin interacts with ISD11 of the NFS1/ISCU complex and multiple mitochondrial chaperones. Hum Mol Genet 16, 929–941, doi:10.1093/hmg/ddm038 (2007).

18 Cai, K., Frederick, R. O., Dashti, H. & Markley, J. L. Architectural Features of Human Mitochondrial Cysteine Desulfurase Complexes from Crosslinking Mass Spectrometry and Small-Angle X-Ray Scattering. Structure 26, 1127–1136 e1124, doi:10.1016/j.str.2018.05.017 (2018).

19 Prischi, F. et al. Structural bases for the interaction of frataxin with the central components of iron-sulphur cluster assembly. Nat Commun 1, 95, doi:10.1038/ncomms1097 (2010).

20 Cai, K., Frederick, R. O., Tonelli, M. & Markley, J. L. Interactions of iron-bound frataxin with ISCU and ferredoxin on the cysteine desulfurase complex leading to Fe-S cluster assembly. J Inorg Biochem 183, 107–116, doi:10.1016/j.jinorgbio.2018.03.007 (2018).

21 Bridwell-Rabb, J., Fox, N. G., Tsai, C. L., Winn, A. M. & Barondeau, D. P. Human frataxin activates Fe-S cluster biosynthesis by facilitating sulfur transfer chemistry. Biochemistry 53, 4904–4913, doi:10.1021/bi500532e (2014).

22 Yoon, H. et al. Frataxin-bypassing Isu1: characterization of the bypass activity in cells and mitochondria. Biochem J 459, 71–81, doi:10.1042/BJ20131273 (2014).

23 Yoon, H. et al. Turning Saccharomyces cerevisiae into a Frataxin-Independent Organism. PLoS Genet 11, e1005135, doi:10.1371/journal.pgen.1005135 (2015).

24 Yan, R. et al. Ferredoxin competes with bacterial frataxin in binding to the desulfurase IscS. J Biol Chem 288, 24777–24787, doi:10.1074/jbc.M113.480327 (2013).

25 Lawrence, N. S. et al. The electrochemical analog of the methylene blue reaction: A novel amperometric approach to the detection of hydrogen sulfide. Electroanal 12, 1453–1460, doi:Doi 10.1002/1521-4109(200012)12:18<1453::Aid-Elan1453>3.0.Co;2-Z (2000).

26 Tsai, C. & Barondeau, D. P. Human frataxin is an allosteric switch that activates the Fe-S cluster biosynthetic complex. Biochemistry 49, 9132–9139, doi:10.1021/bi1013062 (2010).

27 Petoukhov, M. V. et al. New developments in the ATSAS program package for small-angle scattering data analysis. J Appl Crystallogr 45, 342–350, doi:10.1107/S0021889812007662 (2012).

28 Franke, D. & Svergun, D. I. DAMMIF, a program for rapid ab-initio shape determination in small-angle scattering. J Appl Crystallogr 42, 342–346, doi:10.1107/S0021889809000338 (2009).

29 Volkov, V. V. & Svergun, D. I. Uniqueness of ab initio shape determination in small-angle scattering. Journal of Applied Crystallography 36, 860–864, doi:doi:10.1107/S0021889803000268 (2003).

30 Schneidman-Duhovny, D., Hammel, M., Tainer, J. A. & Sali, A. Accurate SAXS profile computation and its assessment by contrast variation experiments. Biophys J 105, 962–974, doi:10.1016/j.bpj.2013.07.020 (2013).

31 Schneidman-Duhovny, D., Hammel, M., Tainer, J. A. & Sali, A. FoXS, FoXSDock and MultiFoXS: Single-state and multi-state structural modeling of proteins and their complexes based on SAXS profiles. Nucleic Acids Res 44, W424–W429, doi:10.1093/nar/gkw389 (2016).

32 Zheng, S. Q. et al. MotionCor2: anisotropic correction of beam-induced motion for improved cryoelectron microscopy. Nat Methods 14, 331–332, doi:10.1038/nmeth.4193 (2017).

33 Zhang, K. Gctf: Real-time CTF determination and correction. J Struct Biol 193, 1–12, doi:10.1016/j.jsb.2015.11.003 (2016).

34 Scheres, S. H. & Chen, S. Prevention of overfitting in cryo-EM structure determination. Nat Methods 9, 853–854, doi:10.1038/nmeth.2115 (2012).

35 Penczek, P. A., Grassucci, R. A. & Frank, J. The ribosome at improved resolution: new techniques for merging and orientation refinement in 3D cryo-electron microscopy of biological particles. Ultramicroscopy 53, 251–270 (1994).

36 Rosenthal, P. B. & Henderson, R. Optimal determination of particle orientation, absolute hand, and contrast loss in single-particle electron cryomicroscopy. J Mol Biol 333, 721–745 (2003).

37 Kucukelbir, A., Sigworth, F. J. & Tagare, H. D. Quantifying the local resolution of cryo-EM density maps. Nat Methods 11, 63–65, doi:10.1038/nmeth.2727 (2014).

38 Arnold, K., Bordoli, L., Kopp, J. & Schwede, T. The SWISS-MODEL workspace: a web-based environment for protein structure homology modelling. Bioinformatics 22, 195–201, doi:10.1093/bioinformatics/bti770 (2006).

39 Pettersen, E. F. et al. UCSF Chimera--a visualization system for exploratory research and analysis. J Comput Chem 25, 1605–1612, doi:10.1002/jcc.20084 (2004).

40 Emsley, P. & Cowtan, K. Coot: model-building tools for molecular graphics. Acta Crystallogr D Biol Crystallogr 60, 2126–2132, doi:10.1107/S0907444904019158 (2004).

41 Adams, P. D. et al. PHENIX: a comprehensive Python-based system for macromolecular structure solution. Acta Crystallogr D Biol Crystallogr 66, 213–221, doi:10.1107/S0907444909052925 (2010).

42 Barad, B. A. et al. EMRinger: side chain-directed model and map validation for 3D cryo-electron microscopy. Nat Methods 12, 943–946, doi:10.1038/nmeth.3541 (2015).

43 Schrodinger, LLC. The PyMOL Molecular Graphics System, Version 1.8 (2015).

