## Supplementary tables & figures for "Structure of the human frataxin-bound iron-sulfur cluster assembly complex provides insight into its activation mechanism"

*Correspondence should be addressed to

**This Supplementary File contains:**

Supplementary Tables S1-S2

Supplementary Figures S1-S9

**Table S1. Data collection, reconstruction, and model refinement statistics**

|  | SDAUF | SDAUF |
| --- | --- | --- |
| **Data collection** |  |  |
| Microscope | Titan Krios | Titan Krios |
| Voltage (keV) | 300 | 300 |
| Nominal magnification | 22,500 x | 22,500 x |
| Exposure navigation | Stage Position | Stage Position |
| Electron exposure (e /Å^2^) | 42 | 42 |
| Dose rate (e/pixel/sec) | 5 | 5 |
| Detector | K2 Summit | K2 Summit |
| Pixel size (Å)* | 0.543 | 0.543 |
| Defocus range (µm) | 1.2 to 2.5 | 1.2 to 2.5 |
| Micrographs Used | 4260 | 4260 |
| Final Refined particles (no.) | 267,153 | 267,153 |
| **Reconstruction** |  |  |
| Symmetry imposed | C2 | C1 |
| Resolution (global) |  |  |
| FSC 0.143 | 3.24 Å | 3.39 Å |
| Applied B-factor (Å^2^) | -178 | -151 |
| **Refinement** |  |  |
| Protein residues  Ligands | 1592  6 |  |
| Map Correlation Coefficient | 0.824 |  |
| R.m.s deviations |  |  |
| Bond lengths (Å) | 0.006 |  |
| Bond angles (°) | 0.646 |  |
| Ramachandran |  |  |
| Outliers | 0.00 % |  |
| Allowed | 7.32 % |  |
| Favored | 92.68 % |  |
| Poor rotamers (%) | 0.00 % |  |
| MolProbity score | 1.81 |  |
| EMRinger score | 4.77 |  |
| Clashscore (all atoms) | 5.71 |  |

*Calibrated pixel size at the detector

**Table S2 Proposed acyl-chain structures and relative abundances of acyl-ACP based on LC-MS**

| **m/z (4’-PP acyl chains)** | **elemental compositions*** | **predicted m/z** | **mass error (mDa)^#^** | **acyl structures^$^** | **acyl-chain length** | **relative abundances (%)^&^** |
| --- | --- | --- | --- | --- | --- | --- |
| 443.295 | C_23_H_42_N_2_O_4_SH^+^ | 443.294 | 1.0 | saturated acyl | C12 | 5.7 |
| 471.324 | C_25_H_46_N_2_O_4_SH^+^ | 471.326 | -2.0 | saturated acyl | C14 | 5.0 |
| 485.304 | C_25_H_44_N_2_O_5_SH^+^ | 485.305 | -1.0 | 3-ketoacyl | C14 | 14.0 |
| 487.320 | C_25_H_46_N_2_O_5_SH^+^ | 487.321 | -1.0 | 3-hydroxyacyl | C14 | 5.1 |
| 497.341 | C_27_H_48_N_2_O_4_SH^+^ | 497.341 | 0.0 | 2-enoylacyl | C16 | 15.5 |
| 499.358 | C_27_H_50_N_2_O_4_SH^+^ | 499.357 | 1.0 | saturated acyl | C16 | 8.5 |
| 513.338 | C_27_H_48_N_2_O_5_SH^+^ | 513.336 | 2.0 | 3-ketoacyl | C16 | 22.6 |
| 525.372 | C_29_H_52_N_2_O_4_SH^+^ | 525.373 | -1.0 | 2-enoylacyl | C18 | 14.9 |
| 541.369 | C_29_H_52_N_2_O_5_SH^+^ | 541.368 | 1.0 | 3-ketoacyl | C18 | 8.7 |

^*^Elemental compositions were derived based on accurate mass measurment.

^#^The mass errors are the differences between the experimental and predicted values expressed in mDa.

^$^Acyl-chain structures were proposed based on elemental compositions and the MS/MS/MS fragments.

^&^Relative abundances were derived based on extracted ion chromatograms of the 6+ holy-ACP molecular ions.

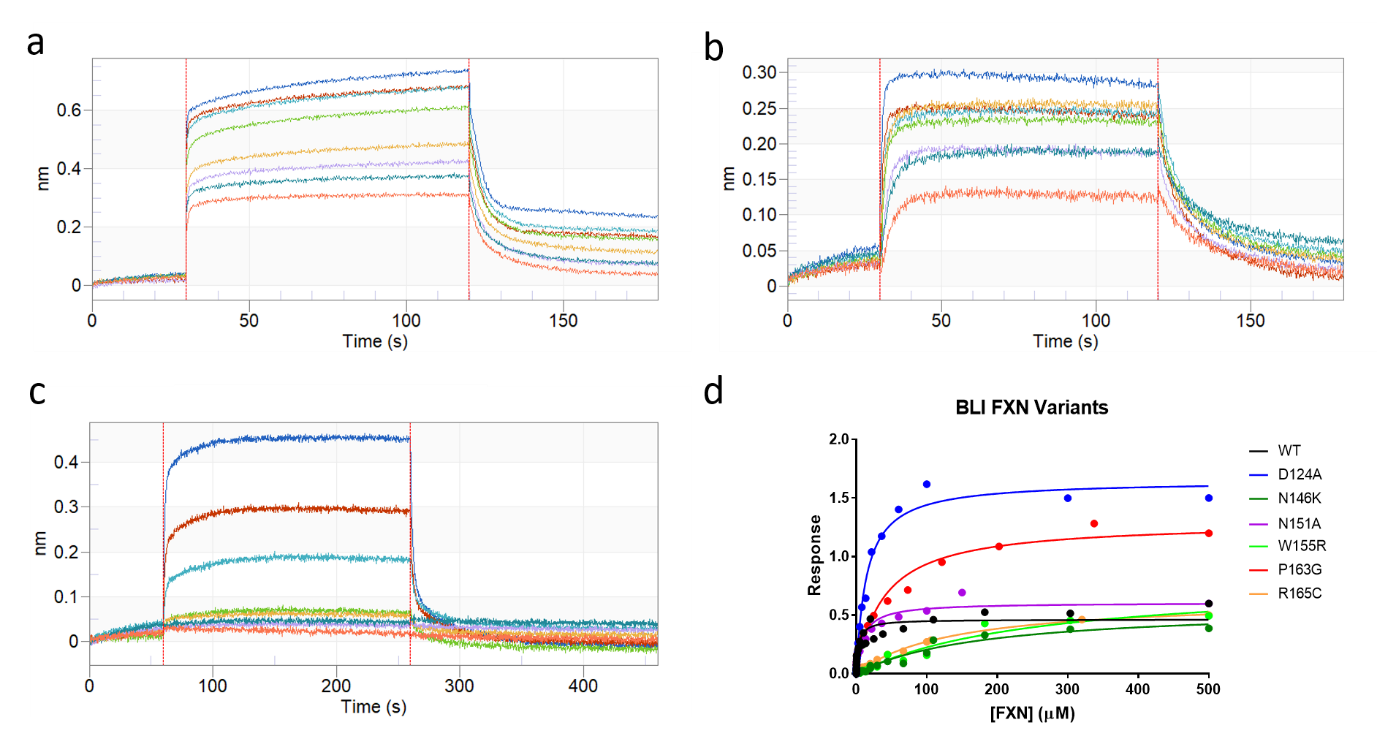

**Figure S1 Bio-layer interferometry of FXN WT and variants.** Raw data curve for FXN(WT) binding to a biotinylated complex of ISCU with SDA_ec_ and titrating various concentrations of FXN ranging from 14-500 μM (**a**), 0.25-8 μM (**b**), and 0-10 μM (**c**). **d.** A plot of response vs. [FXN] (WT and variants) was used for *K_d_* determination using one site-specific binding fit in GraphPad Prism (values reported in Fig. 2e).

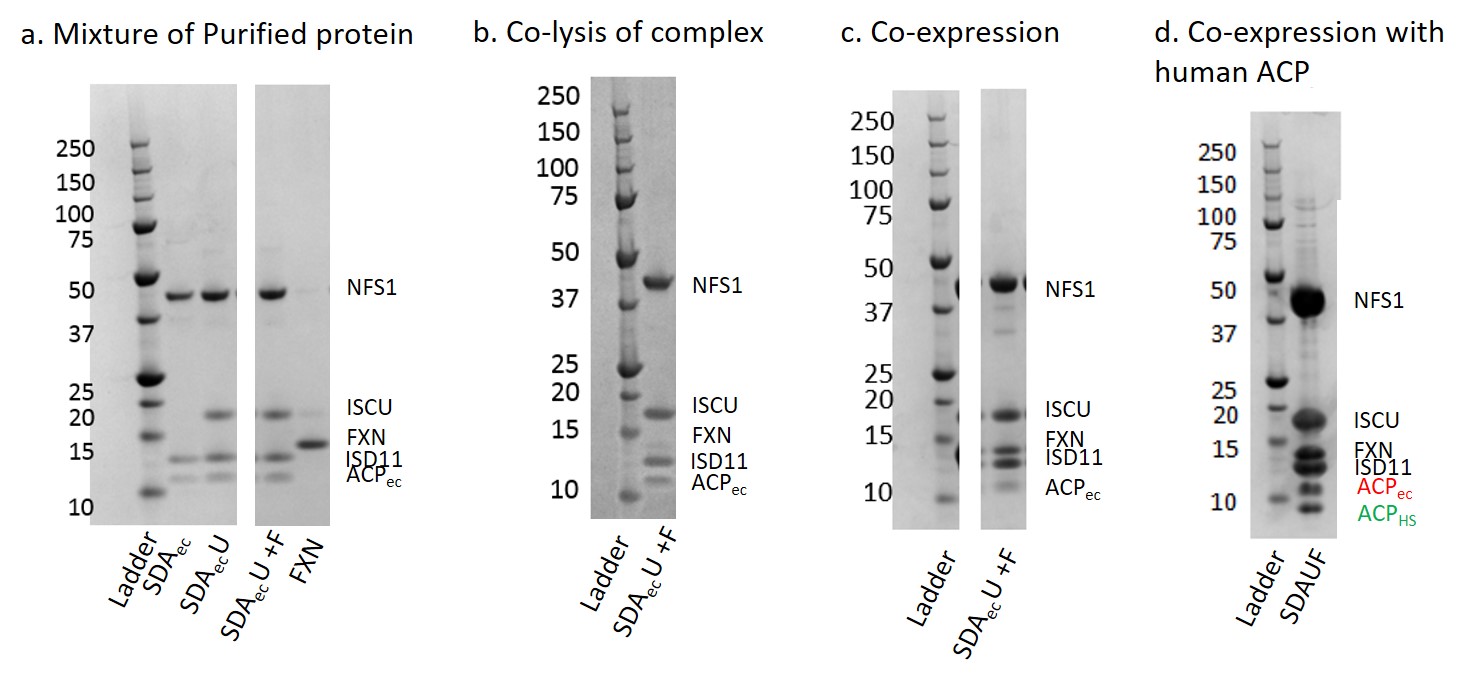

**Figure S2 Different attempts of purifying the SDAUF Complex. a.** Pellets from the expression of the bi-cistronic plasmid (His-ISD11 and NFS1) and tri-cistronic plasmid (His-ISD11, NFS1, and ISCU) were purified and shown to co-purify with *E. coli* ACP (yielding SDA_ec_ and SDA_ec_U, respectively). Purified FXN was added to purified SDA_ec_U complex and run on gel filtration where no FXN binding was observed (SDA_ec_U +F). **b.** Pellets from tri-cistronic expression and FXN expression were co-lysed and purified, showing very little FXN bound. **c.** Co-expression of tri-cistronic plasmid with FXN showed a 5-way complex of SDUF co-purifying with E. coli ACP. **d.** Co-expression of tri-cistronic plasmid with FXN and human ACP showed a 5-way complex of SDAUF with a mixture of both human and *E. coli* ACP.

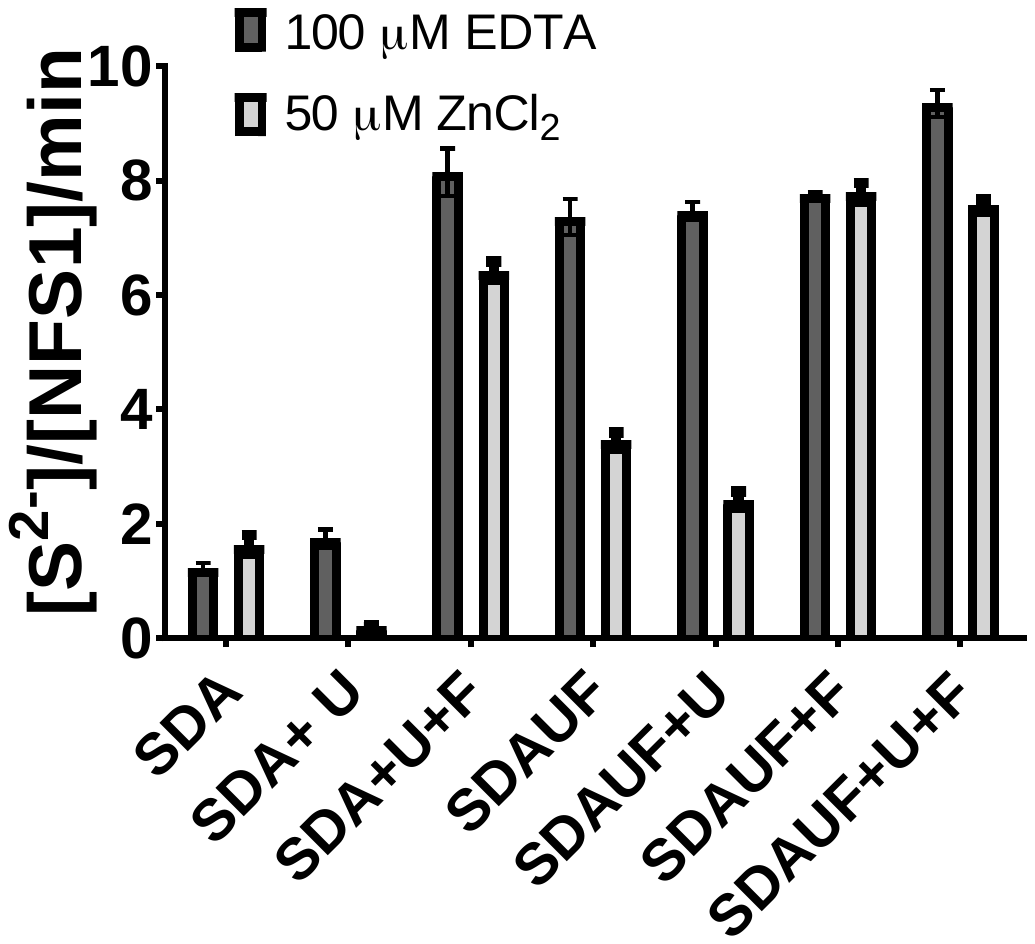
**Figure S3 Activity assay of purified SDAUF complex.** NFS1 activity was measured by the methylene blue assay for individual complexes and combinations with and without Zn^2+^. All complexes co-purified with E. coli ACP. The basal active form of complexes (SDA and SDA+U) could become fully activated when all 5 proteins were added (SDA+U+F, in a ratio of 1:5:5, respectively) regardless of the zinc inhibitor. The isolated complex (SDAUF) showed activity as great as when assayed with all components added separately, indicating that an activated complex was isolated in purification. When zinc was added to the assay, there was still some inhibition for the SDAUF complex, most likely due to FXN not staying fully bound to the complex. Adding additional ISCU (SDAUF+U) further reduced NFS1 activity, as ISCU is the target for zinc inhibition. The addition of more FXN (SDAUF+F or SDAUF+U+F) was able to overcome the zinc inhibition completely.

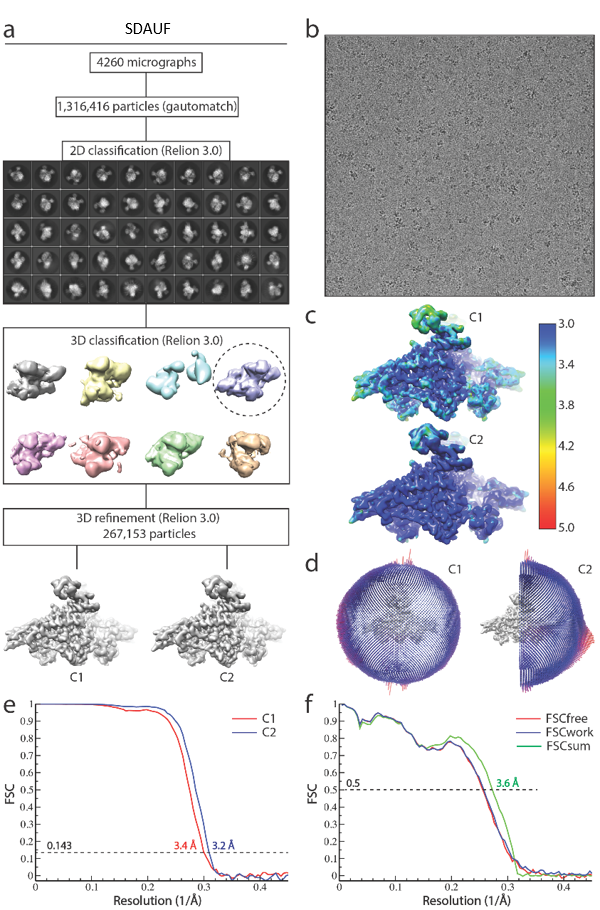

**Figure S4 Cryo-EM analysis of SDAUF complex.** **a.** Flow chart of the cryo-EM data processing procedure. Details can be found in the Methods. **b.** A representative cryo-EM micrograph. **c.** Local resolution of the maps estimated using the ResMap program and colored as indicated. **d.** Angular orientation distribution of the particles used in the final reconstruction. The particle distribution is indicated by different color shades. **e**. Gold standard FSC curves of the structures with or without imposing C2 symmetry. **f.** Model validation. Comparison of the FSC curves between model and half map 1 (work), model and half map 2 (free), and model and full map are plotted in blue, red, and green, respectively.

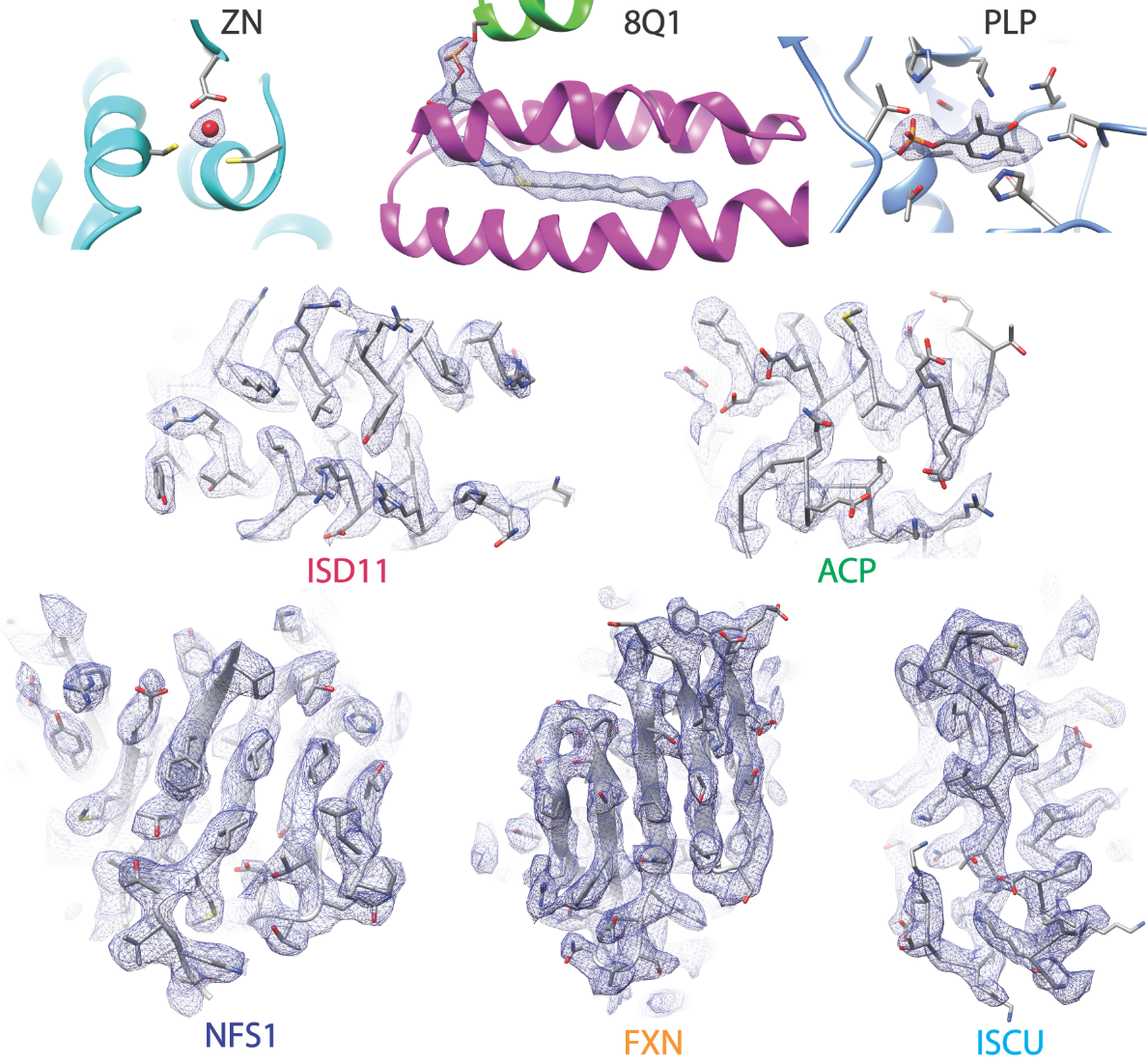

**Figure S5 Fit of the model to the density.** Residues from SDAUF complex (in stick representation) and surrounding electron density maps are shown for exemplary regions. Cryo-EM density is displayed at the contour level 5-6.5 σ around the atoms. Subunits, ligands and maps are specified. Regions with best resolution include NFS1, ISCU, and FXN as well as ISD11. Poorest map quality was observed in the ACP regions.

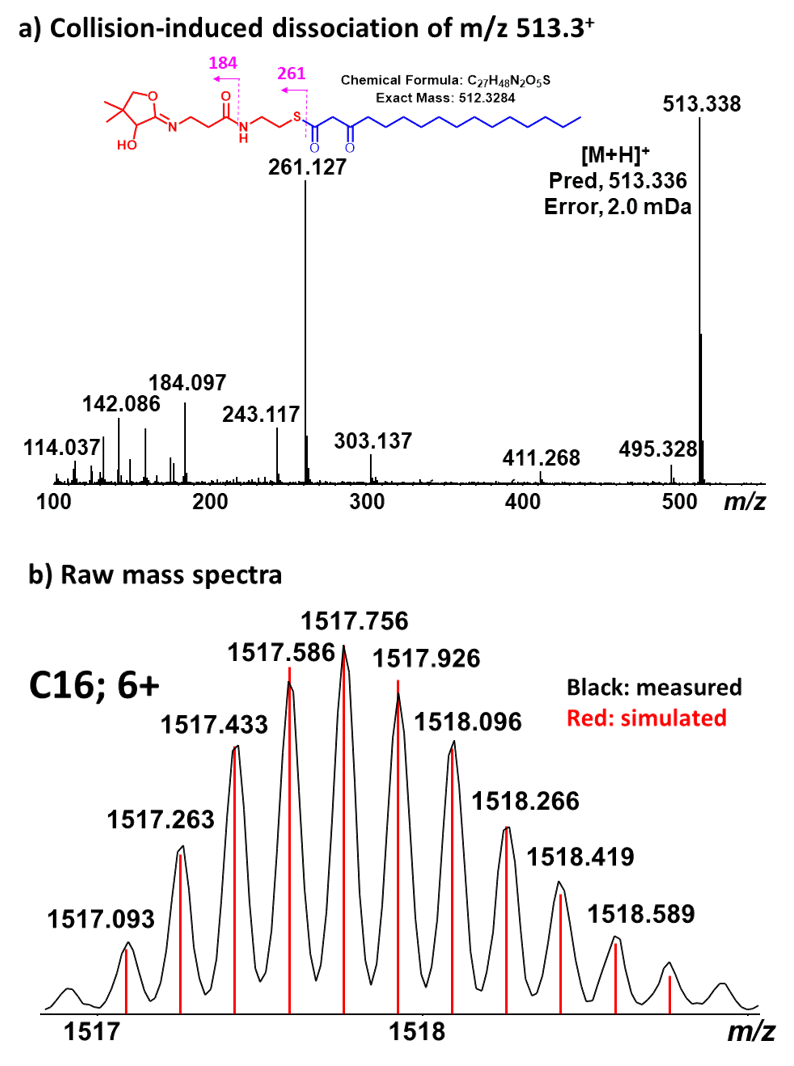

**Figure S6**. Using 3-ketoacyl-ACP (C16) as a representative example to show **a.** MS/MS/MS spectrum for the ejected 4’-PPT conjugated acyl-chains with both parent and fragment ions detected by ppm mass accuracy. Insert shows the proposed structure and fragmentation pathways. **b.** Measured isotopic distribution (black) of the 6+ ion of 3-ketoacyl-ACP (C16) agreed very well with the simulated one (red) by both relative abundances and ppm mass accuracy for all the detectable isotopes strongly supporting the proposed structure.

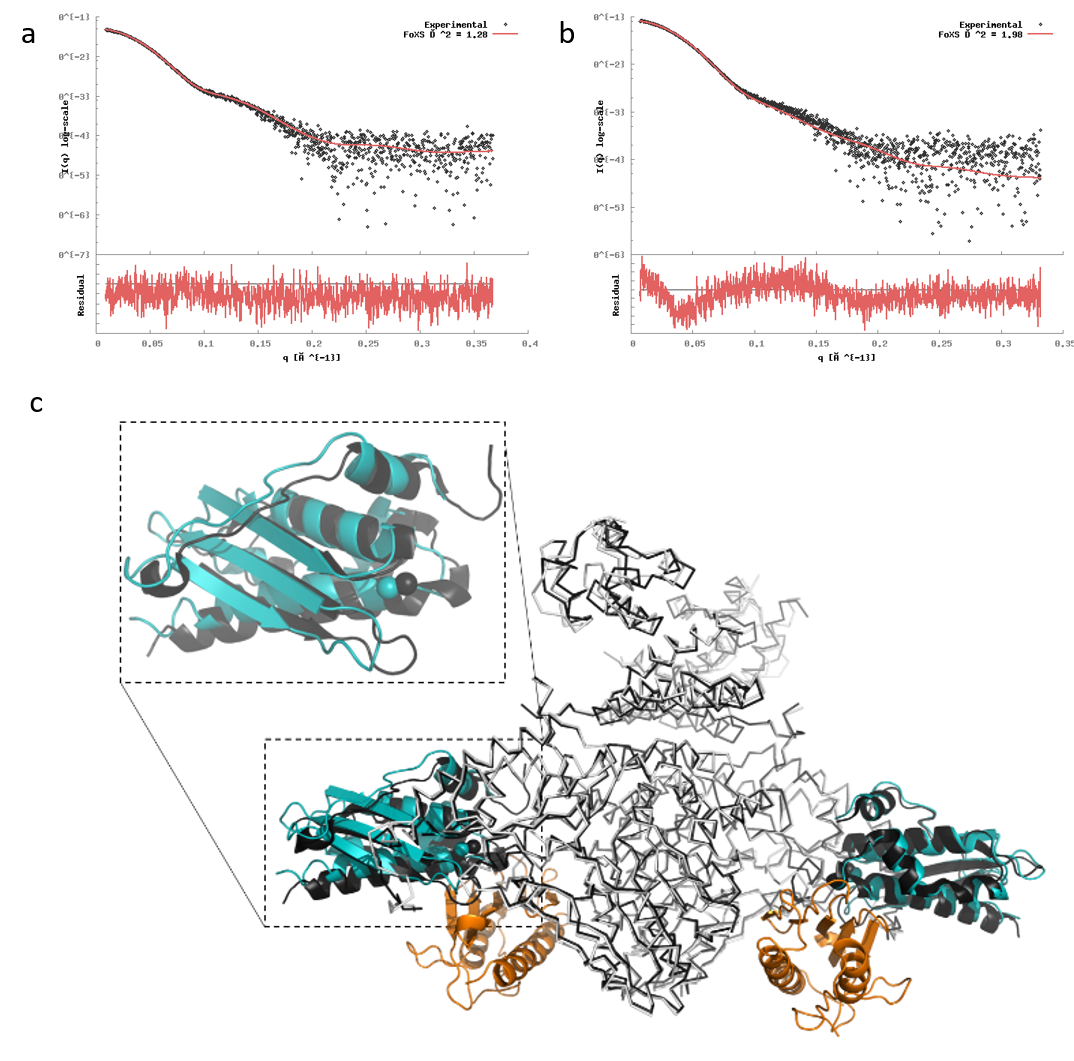

**Figure S7** SAXS analysis and structural superposition of complexes. **a.** Scattering data from a sample of SDAU was collected (black points) and fit to the theoretical SAXS profile back-calculated from SDAU-Zn^2+^ structure (5WLW)(red line) with χ^2^=1.28. **b.** Scattering data from a sample of SDAUF used in cryo-EM was collected (black points) and fit to the theoretical SAXS profile back-calculated from the SDAUF-Zn^2+^ cryo-EM structure (red line) with χ^2^=1.98. **c**. Superimposition of Cα chains from SDAUF-Zn^2+^ structure (this study; NFS1, ISD11, and ACP in white, ISCU in cyan, and FXN in orange) with crystal structures of SDAU-Zn^2+^ (5WLW; all subunits in black) showing a displacement of ISCU within the complex core upon FXN binding.

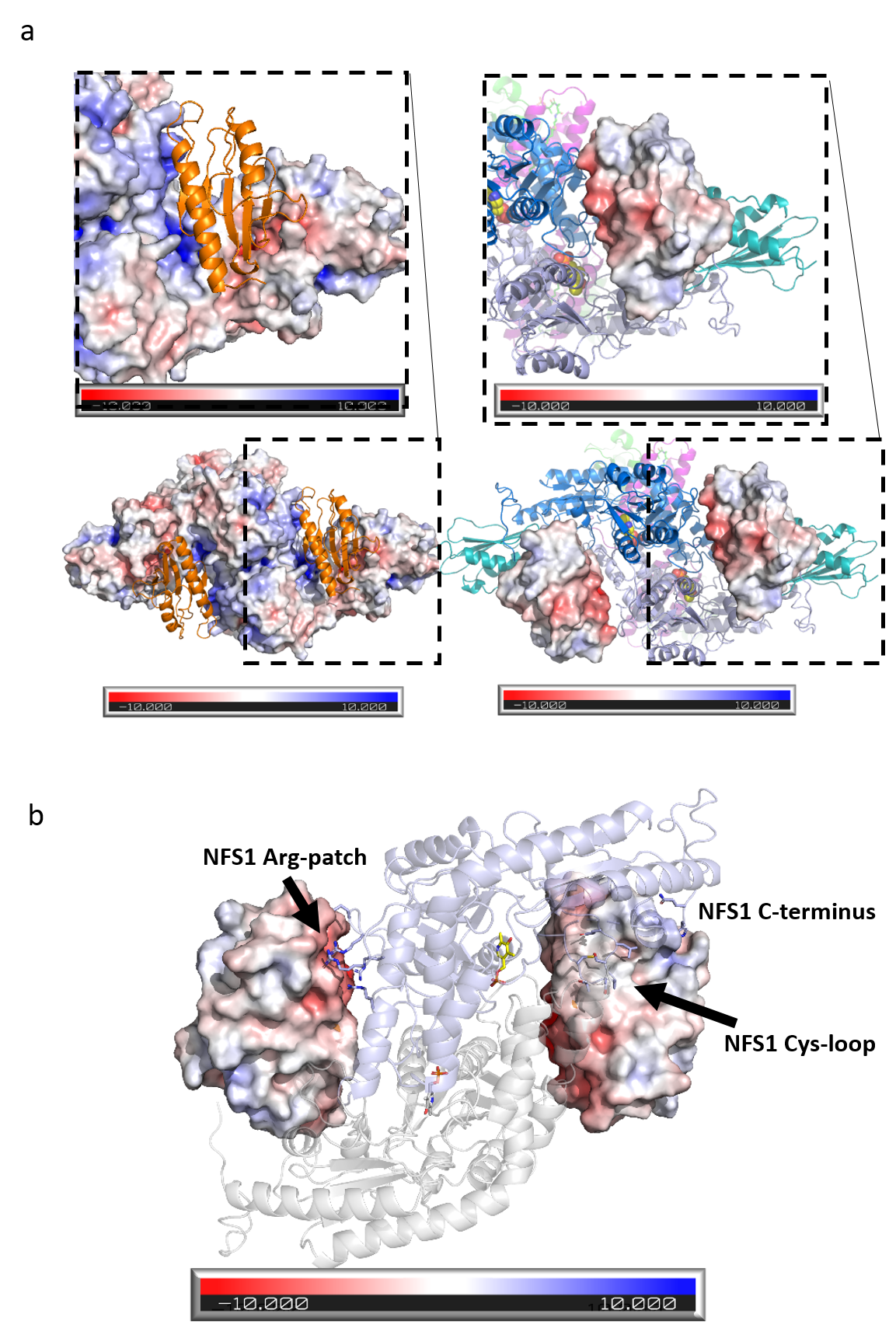

**Figure S8** **a**. Left: Surface representation of SDAU (from the SDAUF-Zn^2+^ structure), colored by electrostatic potential, shows a positively charged surface where FXN (orange) interacts. Right: Surface representation of FXN shows its negative surface that interacts with the positively charged surface on NFS1. **b**. View down the 2-fold axis of NFS1 homodimer (slate/gray), showing interactions with both FXN protomers (surface representation) of the complex per NFS1 . Each NFS1 uses its Cys-loop and C-terminus to contact one FXN protomer, and uses its Arg-patch on the other side of NFS1 to contact the other FXN protomer.

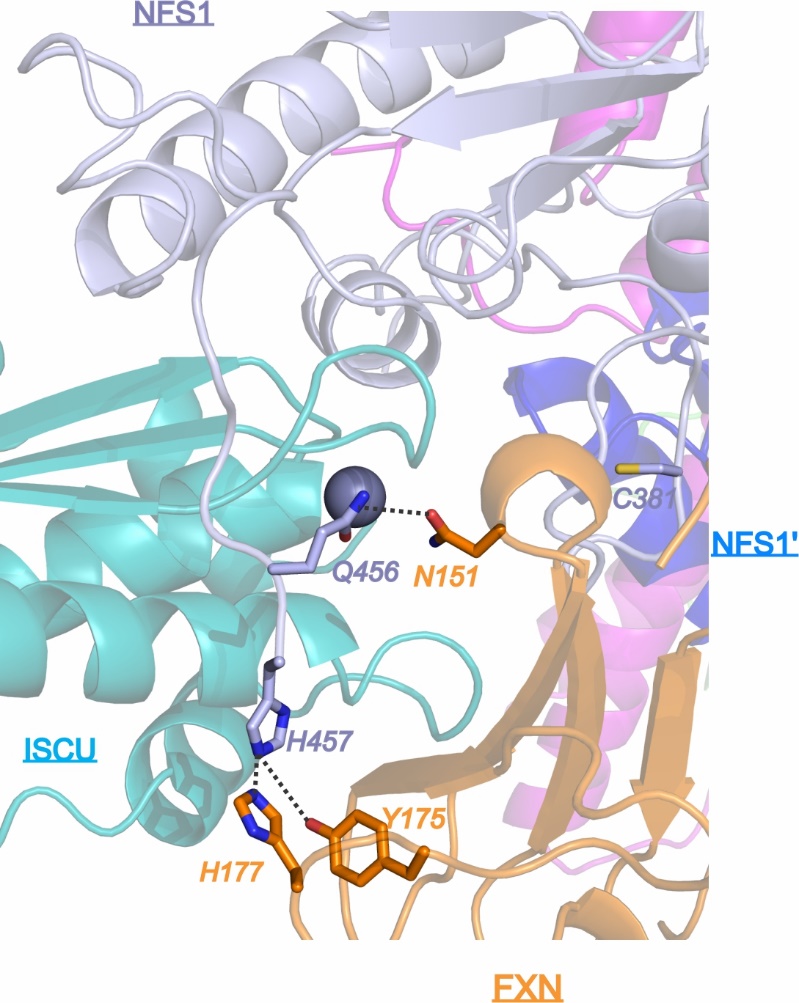

**Figure S9** NFS1 C-terminus (slate) wraps around ISCU (cyan), with the NFS1 terminal residues (Gln456 and His457) interacting directly with FXN Asn151, Tyr175, His177 (orange).
